# PowerFit: An updated framework for fast rigid-body fitting into cryo-EM density maps

**DOI:** 10.64898/2026.09.29.755289

**Authors:** Anna Engel, Stefan Verhoeven, Bart Schilperoort, Rodrigo Vargas Honorato, Alexandre M.J.J. Bonvin

**Affiliations:** Bijvoet Centre for Biomolecular Research, Faculty of Science – Chemistry, Utrecht University, Utrecht, the Netherlands; The Netherlands eScience Center, Amsterdam, the Netherlands

**Keywords:** Cryo-EM, Exhaustive sampling, GPU acceleration

## Abstract

Cryo-electron microscopy (cryo-EM) has become a central method for determining structures of molecular assemblies, requiring reliable tools to interpret the density maps. Despite recent advances in AI-driven model building, at intermediate-to-low resolution (4-15 Å) accurate rigid-body fitting of existing atomic structures into cryo-EM maps remains an essential step in the modelling process. Here we present an updated version of our software package PowerFit. It performs an exhaustive translational and rotational search against the cryo-EM density map, implementing sensitive correlation functions. The updated release preserves the fitting accuracy and sensitivity of the original method while substantially improving the computational performance through modernised CPU and GPU implementations. In addition, to facilitate the interpretation of results, it provides an HTML report with interactive 3D visualisation using MolViewSpec. The same HTML report is used to generate the result page when using PowerFit via its web service. The report enables immediate inspection and validation of the results. The software’s accessibility has been enhanced by updating it from Python 2 to Python 3, enabling straightforward installation via the Python Package Index (PyPI) and simple integration into computational pipelines. PowerFit is freely available at https://github.com/haddocking/powerfit, as a web service at https://wenmr.science.uu.nl/powerfit or in JupyterLite directly in a web browser at https://www.bonvinlab.org/powerfit/jupyterlite.html.

**Highlights:**

- Accurate rigid-body fitting of existing atomic models remains an essential step in interpreting density maps at intermediate-to-low resolution.
- New, updated, much faster version of PowerFit preserving fitting accuracy and sensitivity.
- Modernised CPU and GPU implementations to improve performance.
- Interactive HTML report with 3D visualisation and validation of fitting results.
- Accessible through simplified PyPI installation, as a web service or in JupyterLite directly in a web browser.

**Graphical Abstract:** 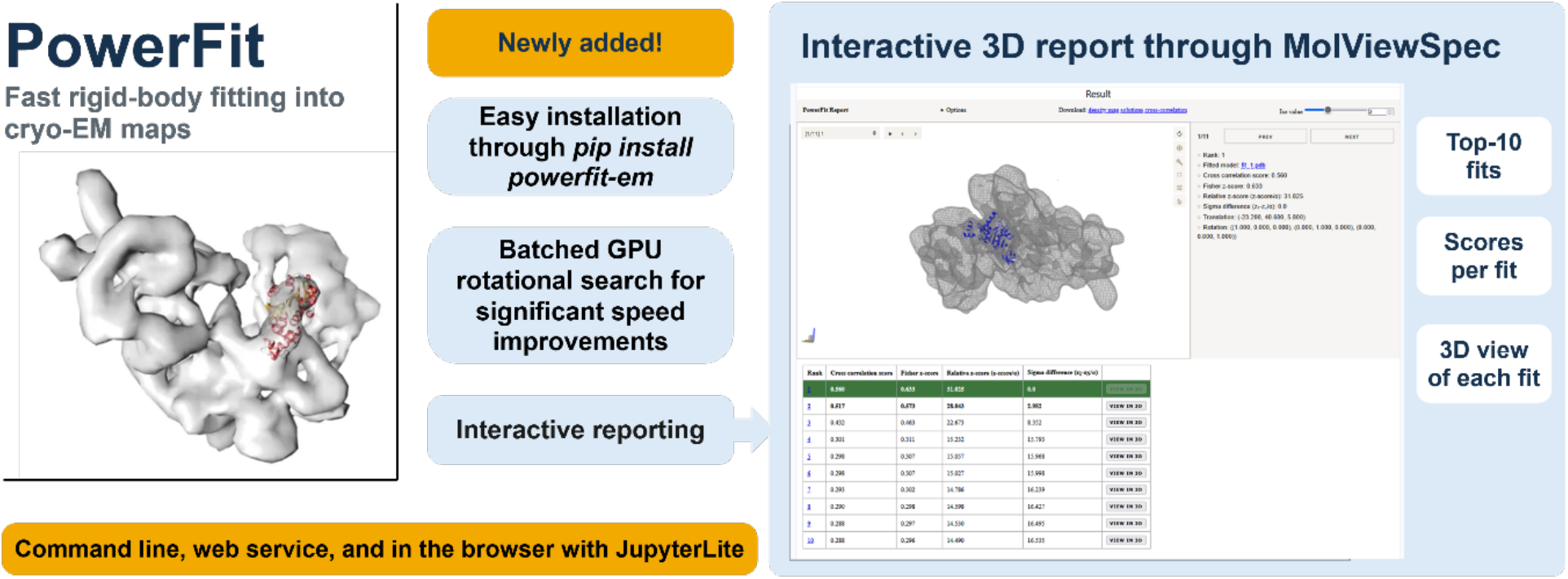

## Introduction

Cryo-Electron Microscopy (Cryo-EM) has become an increasingly important tool for studying protein structures, due to improvements in imaging software and hardware [1,2]. Cryo-EM can solve structures that could not previously be solved by Nuclear Magnetic Resonance (NMR) or X-ray Crystallography (X-ray) due to their size, flexibility or difficulties in purification and crystallisation [3,4]. The output of a Cryo-EM experiment is not an image but a density map representing the three-dimensional structure of the biomolecular system studied at a given resolution. Reliable tools are needed to accurately interpret these density maps and visualise the interactions in biomolecular complexes.

Different tools are available to interpret a 3D density map, depending on its resolution. When a density map has a high resolution (<4.5 Å), the protein backbone can be traced through the density directly, and the structure built from bottom up [5–10]. Densities with a resolution lower than 4.5 Å can no longer be traced this way, so rigid-body fitting tools are needed to interpret them [11]. They require a 3D structural template. This template can be experimentally solved structures available in the Protein Data Bank (PDB) [12] or structural models. More recently, templates can also be directly generated from sequence using deep learning methods such as AlphaFold [13,14]. Once an appropriate 3D template has been selected or generated, a rigid-body fitting tool can be used to place it into the 3D density map by sampling all possible translations and rotations and finding the best-fitting models using some correlation function that measures the fit between the model and the density at a given position [15–19].

Multiple rigid-body fitting tools to place templates into densities have been developed, including Situs [15], ADP_EM [20], the dock-in-map tool in Phenix [21], fit-in-map in ChimeraX [22], TEMPy [23], and VESPER [24]. An additional tool is PowerFit, developed in our lab [25]. PowerFit systematically rotates and translates the template to find possible placements in the density map. It combines a local cross-correlation (LCC) score [15], a core-weighted (CW) cross-correlation score[16], and a Laplace pre-filter [17] for highly sensitive and accurate fitting. It uses the Fast Fourier Transform (FFT) [17] algorithm to accelerate the translational search. In addition, to further speed-up the search, it automatically trims and resamples the density.

When PowerFit was originally published, it was available both as CPU- and GPU-accelerated code, from the command line or as a web service [26]. It was among the fastest rigid-body fitting tools at that time. However, this old PowerFit version, written in Python 2, could not keep pace with newer hardware. The software was still available through a web service running on old hardware [26] and through the SBGrid software installation [27]. Because of broken dependencies, GPU acceleration was no longer possible.

We present here an updated version of our PowerFit package [28], ported to Python 3 with modernised CPU and GPU implementations. It is available on the Python Package Index (PyPI) for easy installation and integration into workflows. It also facilitates the interpretation of the results by providing an HTML report with interactive 3D visualisations through the MolViewSpec viewer [29]. This modern implementation maintains the original accuracy and sensitivity while significantly increasing the computational speed. Compared to current state-of-the-art fitting tools, PowerFit remains among the fastest and most accurate rigid-body EM-fitting software packages to date.

## Materials and Methods

### CPU and GPU modernisation and optimisation

In the updated version of PowerFit, the exhaustive translational and rotational search against the EM density has been modernised and significantly optimised for speed. It now runs on Python 3 and is compatible with Python 3.11 or later. It can be easily installed through PyPI (`pip install powerfit-em`). The CPU backend has been modernised by rewriting the older C/Cython code into Rust. The gpyfft library used to offload calculations to the GPU has been replaced by pyvkFFT [30], which supports both OpenCL and CUDA. Lastly, it now samples rotations in batches on the GPU instead of single rotations, which significantly increases the computational speed.

### Benchmarking

Benchmarking was performed with the PowerFit 6.0.1 release. For the speed and accuracy comparison with the old PowerFit implementation, we used version 2.0.0 for the CPU and 3.0.2 for the GPU. The latter is the first functional release after moving to Python 3 but before optimisation. For the comparison, we used the Cis, Trans and GroES subunits from the GroEL/GroES complex (1GRU) with its associated EM density (EMD-1046) [31] with both fine (4.71°) and coarse (20.81°) rotational sampling angles as described in our previous study [25]. The rotational sampling is based on the optimal rotational angles originally developed by Karney for solid state NMR [32]. We ran GPU timing comparisons on an NVIDIA GeForce GTX 1080 Ti (GPU1) and an NVIDIA RTX 6000 Ada Generation (GPU2). CPU timing comparisons were done with either 1 or 8 threads on an AMD EPYC 7451 24-core.

For measuring the accuracy of the fit with respect to the original 1GRU model (a rigid-body fitted model) we calculated the positional Root Mean Square Deviations (RMSD) using ChimeraX version 1.11 [22]. For the comparison to VESPER [24] we used version 20210908. We ran the computations with PowerFit version 2.0.0, ChimeraX, and VESPER through the SBGrid software environment [27] which provides curated, version-controlled structural biology applications and reproducible execution environments across platforms. Before fitting, we randomly rotated and translated the starting templates using the in-house script provided in the Zenodo dataset [33]. For fitting with VESPER, we first had to simulate maps from the protein templates using ChimeraX and then used those in VESPER as VESPER cannot work directly from PDB files. We then applied the translation and rotation values to the protein templates to generate the fitted model and calculate the RMSD with the original deposited template. Rotational sampling intervals for this comparison were set per target, as listed in Table 2 (10° for the GroEL/GroES subunits and 2YKR chain W [34], 5° for 3ZPZ chain C [35])

All templates, scripts and results can be found on Zenodo (https://zenodo.org/records/22920646).

## Results and Discussion

### Maintaining accuracy and sensitivity

To demonstrate that the code optimisations did not affect PowerFit’s accuracy and sensitivity, we compared the performance of the new version (release 6.0.1) with the original version 2.0.0 (Table S1). For scoring, we used the core-weighted Laplace pre-filter local cross-correlation score (L-CW-LCC), which was previously shown to be the most effective scoring method for PowerFit [25]. We also compared the default trimming and resampling parameters to a run with a full map. For our comparison, we used the Cis, Trans and GroES subunits from the GroEL/GroES complex (1GRU) with its associated EM density (EMD-1046) [31] and used a fine rotational sampling interval of 4.71°, as in our previous study (Figure S1). We fitted the Cis and Trans subunits separately and the GroES ring as a whole. To validate the results, we calculated the RMSD between the PowerFit-calculated fits and the 1GRU reference structure. For the Cis and Trans subunits, we report the average RMSD over the top 7 fits, as the system has 7-fold symmetry. The modernised implementation of PowerFit can fit the GroES subunit, and the seven Cis and Trans subunits into the density with identical RMSD values in both the default trimmed and resampled map and a full map. This shows that sensitivity and accuracy have been maintained across release versions.

### GPU speed improvements

The major improvement efforts for PowerFit focused on optimising the GPU implementation (for CPU timings refer to Table S2). Table 1 compares the GPU speed of release version 6.0.1 to version 3.0.2 (unoptimized GPU version) while fitting the 1GRU subunits into the density.

**Table 1.** GPU speed performance of PowerFit version 3.0.2 versus 6.0.1 on the GroEL/GroES complex sampling at a coarse (20.81°) and fine (4.71°) rotational sampling interval with core-weighted Laplace pre-filter local cross-correlation (L-CW-LCC) scoring. Calculations were performed on an NVIDIA GeForce GTX 1080 TI (GPU1) and an NVIDIA RTX 6000 ADA Generation (GPU2).

| PowerFit timings of the GPU runs (s) |  |  |
| --- | --- | --- |
| Coarse sampling (20.81°) |  |  |
| PowerFit Version | 3.0.2 | 6.0.1 |
| GPU |  |  |
| GPU1 | 10.9 | 6.3 |
| GPU2 | 7.1 | 4.6 |
| Fine sampling (4.71°) |  |  |
| PowerFit Version | 3.0.2 | 6.0.1 |
| GPU |  |  |
| GPU1 | 642.3 | 12.0 |
| GPU2 | 352.3 | 4.8 |

We tested the implementation with two different rotational sampling intervals: a coarse (20.81°) and a fine (4.71°) interval, with the other parameters at default. We ran the speed tests on two GPUs: an NVIDIA GeForce GTX 1080 Ti and an NVIDIA RTX 6000 Ada Generation (GPU1 and GPU2, respectively). GPU optimisation in version 6.0.1 improves the coarse rotations by factors 2.1 and 1.6 on GPU1 and GPU2, respectively. For fine rotational sampling this increases to 53.5 and 73.8. On the NVIDIA RTX 6000 Ada (GPU2) card almost no time difference is observed between coarse and fine rotational sampling. This very significant speed improvement is due mainly to the implementation of batched rotational sampling on the GPU. In the old PowerFit version, only one rotation was sampled at a time, whereas in the new version multiple rotations can be calculated at the same time. A four-fold increase in rotational sampling between fine and coarse searches corresponds to 16 times more rotations sampled. The new GPU implementation thus shows the greatest improvement when using finer rotational sampling.

In the previous PowerFit publication, it was recommended to use trimming options for an initial search, since sampling a full map was expensive. With this new GPU implementation, sampling a full map with fine rotational sampling is possible within an acceptable time period, making initial coarse fitting redundant.

### Fitting performance comparison

The original PowerFit version was compared to another GPU-accelerated rigid-body fitting tool gEMfitter [36]. To the best of our knowledge, no other GPU-accelerated tool has been published since the original PowerFit release. Therefore, we compared the performance of PowerFit against VESPER (CPU-only implementation) [24] for 5 different use cases. We chose VESPER as it also allows changing the rotational sampling interval. We fitted here, again, the individual subunits from 1GRU in its associated density map, chain W from 2YKR in EMD-1884 [34], and chain C from 3ZPZ in EMD-2325 [35] (Table 2, Figure S2).

**Table 2.** Performance and timing comparison of PowerFit versus VESPER, using 10 threads on an AMD EPYC 745 24-core CPU. For comparison PowerFit GPU timings on a NVIDIA RTX 6000 ADA card are provided. For both VESPER and PowerFit, default settings were used. The rotational sampling interval was set per case. When no confident fits were returned, RMSDs are not reported.

| Chain | Rotational sampling interval (°) | Resolution (Å) | Top PowerFit | RMSD (Å)* | Time (s) | GPU Time (s) | Top VESPER | RMSD (Å)* | Time (s) |
| --- | --- | --- | --- | --- | --- | --- | --- | --- | --- |
| 1GRU Cis | 10 | 23.5 | 1-7 | 5.76 | 5.5 | 2.3 | / | / | 7.5 |
| 1GRU Trans | 10 | 23.5 | 1-7 | 6.53 | 5.4 | 2.1 | / | / | 7.4 |
| 1GRU GroES | 10 | 23.5 | 1 | 5.77 | 5.5 | 2.1 | / | / | 7.4 |
| 2YKR chain W | 10 | 9.8 | 1 | 3.17 | 94.3 | 9.7 | / | / | 19.7 |
| 3ZPZ chain C | 5 | 8.9 | 1-7 | 2.02 | 292.8 | 37.7 | / | / | 34.0 |

\*. RMSDs were calculated against the originally deposited templates, which were themselves obtained by rigid-body fitting into the density.

We ran the searches on an AMD EPYC 7451 24-core CPU using 10 threads and maintained default settings. As shown in Table 2 and Figure S2, PowerFit fits the chains in the correct position, with speeds faster than VESPER at low resolution but slower as resolution and rotational sampling interval increase. This difference however disappears using the GPU version of PowerFit, which is as fast as or faster than VESPER. PowerFit performs the correlation search on a voxel grid whose spacing matches the input map resolution. VESPER reduces the map resolution to a coarse set of unit vectors for a global search, then refines only the top-ranked poses in finer detail. The result is that PowerFit is faster at low resolution, but becomes slower as the resolution increases

PowerFit accuracy, however, is better than VESPER in all cases. VESPER uses as significance cutoff a fit’s Z-score of 10 or higher. In all tested cases, the VESPER Z-score was below this cutoff. Therefore, VESPER could not reliably fit any of the templates into the densities. As such no RMSD values are reported. For PowerFit, on all five cases, confident fits are obtained for all subunits based on the Fisher z-score and the confidence interval between fits, which determines whether subsequent fits besides the top-ranked one are also significant [37]. With this metric, all seven subunits of 1GRU Cis, 1GRU Trans, and 3ZPZ chain C could be reliably fitted (Figure S2). This is further validated by the calculated RMSD values between the PowerFit-generated fits and the originally fitted models.

In addition to its better performance, another major advantage of PowerFit over VESPER is that it directly fits the protein template into the density, whereas VESPER first requires manually transforming the protein template into a simulated density map at the same resolution as the target map. VESPER then outputs translation and rotation values, which must be applied to the input protein template to generate the fitted models. So for direct visualisation, PowerFit is easier to use.

### Results reporting

In addition to the code optimisations, we also improved the visual reporting of PowerFit results. Where previously only an output text file with scores was generated, and static images on the web service, we now provide interactive 3D reporting through MolViewSpec (Figure 1A). This HTML report can be generated using the CLI version and is also available on our web service. It provides, users with the top 10 fitted models and their associated score and translation/rotation matrices. Users can also adjust the density contour level to improve visibility. Lastly, all fits can be displayed at once.

**Figure 1.**
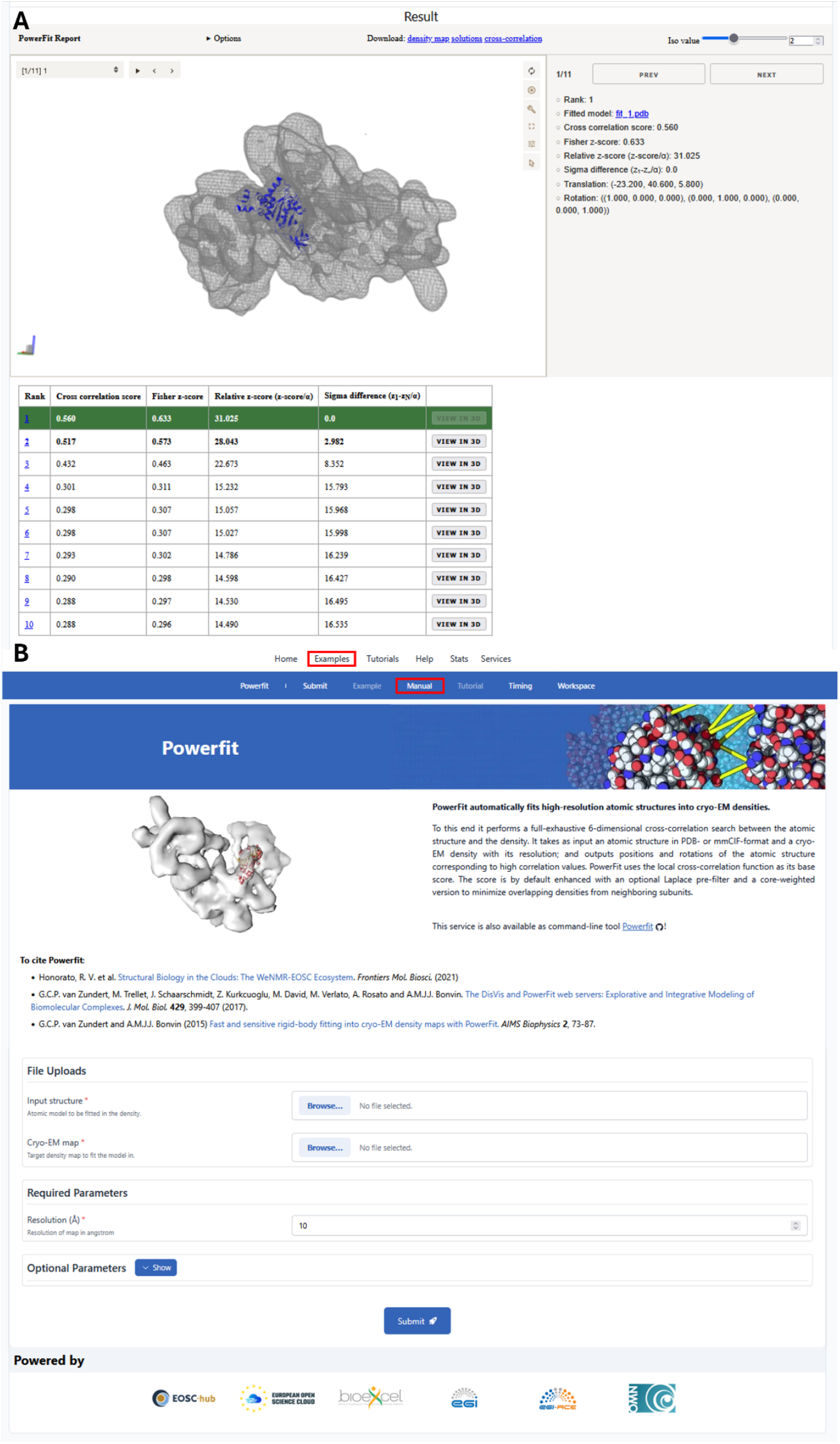
The PowerFit user interfaces. (**A**) Reporting page generated after a successful PowerFit run. (**B**) Submission page of the PowerFit web service (https://wenmr.science.uu.nl/powerfit). Red boxes indicate clickable areas linking to examples and the online manual.

### PowerFit web service

In addition to its command-line version, PowerFit is also available as a web service at https://wenmr.science.uu.nl/powerfit (Figure 1B). The web service runs the same underlying PowerFit code and generates the same interactive HTML report as the command line version, allowing users to submit a job and inspect results without any local installation. To help new users get started, examples are provided (https://wenmr.science.uu.nl/examples). A manual describing both command-line and web service usage is also available at https://bonvinlab.org/powerfit. These are also directly accessible when on the submission page (Figure 1B, red boxes). The new PowerFit web service has run to date (27-09-2026) 435 jobs (https://wenmr.science.uu.nl/stats#jobs).

## Conclusions

We have presented the modernised and optimised version of our open-source software package PowerFit, which maintains the original fitting accuracy and sensitivity. This new implementation offers significant improvements in speed, especially with the new GPU implementation, which has support for AMD and Intel devices. Its interactive, HTML-based reports provide a user-friendly analysis interface. It is easy to install via PyPI, which facilitates its integration into workflows, and available for various operating systems (Windows, Linux, and MacOSX). Finally, to facilitate its use by less experienced users or those with limited computational resource, it is also available both as a web service (https://www.wenmr.science.uu.nl/powerfit) and a JupyterLite notebook (https://www.bonvinlab.org/powerfit/jupyterlite.html) that can run directly in your web browser.

## CRediT authorship contribution statement

**Anna Engel:** Conceptualisation, Data curation, Formal analysis, Investigation, Methodology, Project administration, Validation, Visualisation, Writing – original draft, Writing – review & editing. **Stefan Verhoeven:** Data curation, Methodology, Project administration, Software, Writing – review & editing. **Bart Schilperoort:** Data curation, Methodology, Software, Writing – review & editing. **Rodrigo Vargas Honorato:** Data curation, Methodology, Software, Writing – review & editing. **Alexandre Bonvin:** Conceptualisation, Funding acquisition, Project administration, Resources, Writing – review & editing.

## Funding

This work was supported by the Dutch Research Council (NWO - OCENW.M.22.349) and the Netherlands eScience Center (NLESC.OEC.2024.3147).

## Data and Software Availability

PowerFit is freely available with a permissive Apache2 license for Windows, Linux, and Mac at https://github.com/haddocking/powerfit or as a web service at https://wenmr.science.uu.nl/powerfit. A user manual describing both its usage at the command line and as a web service, together with example input and output files, is provided at https://bonvinlab.org/powerfit. Finally, PowerFit can also be directly run from a web browser through native JupyterLite integration at https://www.bonvinlab.org/powerfit/jupyterlite.html. All templates, scripts and data described in this paper can be found at : https://zenodo.org/records/22920646.

## Declaration of Competing Interest

The authors have nothing to declare.

## Notes

https://zenodo.org/records/22920646

https://github.com/haddocking/powerfit

https://wenmr.science.uu.nl/powerfit

https://bonvinlab.org/powerfit

### Competing Interest Statement

The authors have declared no competing interest.

