## Supplementary material for "PowerFit: An updated framework for fast rigid-body fitting into cryo-EM density maps"

**Table S1**. Fitting performance of PowerFit version 2.0.0 versus 6.0.1 on the GroEL/GroES complex (1GRU into EMD-1046) using the core-weighted Laplace pre-filter local cross-correlation (L-CW-LCC) score. Cis and Trans RMSD values are averaged of the top-7 fits.

| **Average RMSD values of the fitted subunits (Å)*** | | |
|---|---|---|
| Trimmed and resampled | | |
| **PowerFit Version**<br>**1GRU Domain** | **2.0.0** | **6.0.1** |
| Cis | 5.57 | 5.57 |
| Trans | 6.33 | 6.33 |
| GroES | 7.52 | 7.52 |
| Full map | | |
| **PowerFit Version**<br>**1GRU Domain** | **2.0.0** | **6.0.1** |
| Cis | 4.10 | 4.10 |
| Trans | 2.98 | 2.98 |
| GroES | 4.79 | 4.79 |

*. RMSDs were calculated against the originally deposited templates, which were themselves obtained by rigid-body fitting into the density

**Table S2.** CPU speed performance of PowerFit version 3.0.2 versus 6.0.1 on the GroEL/GroES complex (1GRU into EMD-1046) sampling at a coarse (20.81°) and fine (4.71°) rotational sampling interval using the core-weighted Laplace pre-filter local cross-correlation (L-CW-LCC) score. Calculations were performed on an AMD EPYC 7451 24-core Processor using 1 or 8 threads.

| **PowerFit timings of the CPU runs (s)** | | |
|---|---|---|
| **Coarse sampling** (20.81°) | | |
| **PowerFit Version**<br>**#CPU threads** | **2.0.0** | **6.0.1** |
| 1 | 00:11.8 | 00:10.5 |
| 8 | 00:05.6 | 00:07.5 |
| **Fine sampling** (4.71°) | | |
| **PowerFit Version**<br>**#CPU threads** | **3.0.2** | **6.0.1** |
| 1 | 744.2 | 733.2 |
| 8 | 99.7 | 98.2 |

**Figure S1**. The density (EMD-1046) and template model (1GRU) for the GroEL/GroES complex.

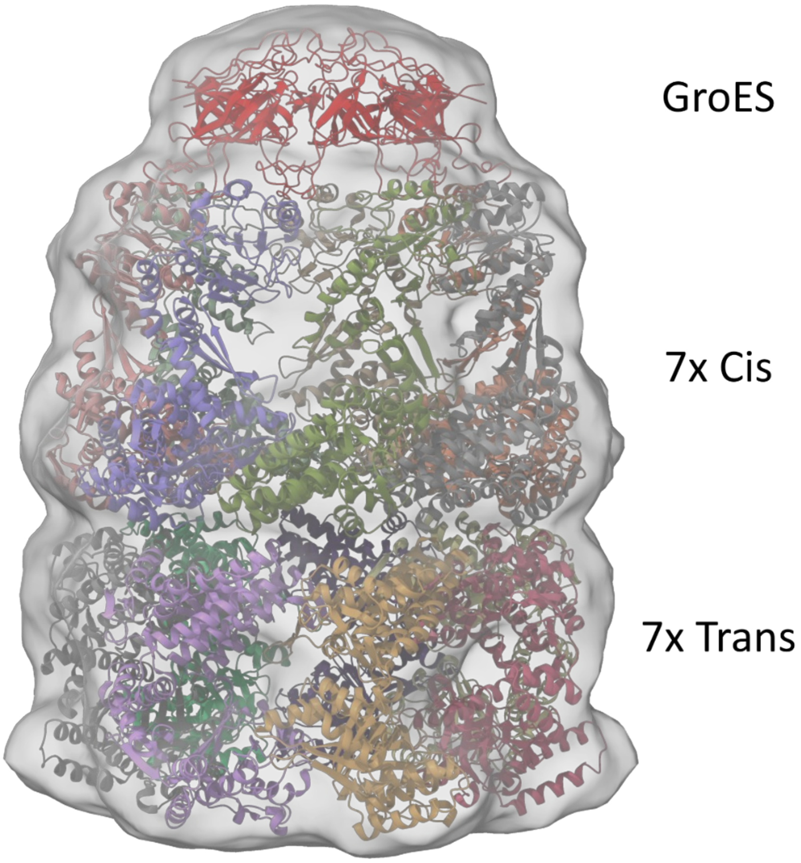

**Figure S2**. ChimeraX images of subunits fitted with PowerFit. (A) Cis, Trans, and GroES subunits (EMD-1046), (B) 2YKR chain W (EMD-1884), (C) 3ZPZ chain C (EMD-2325).

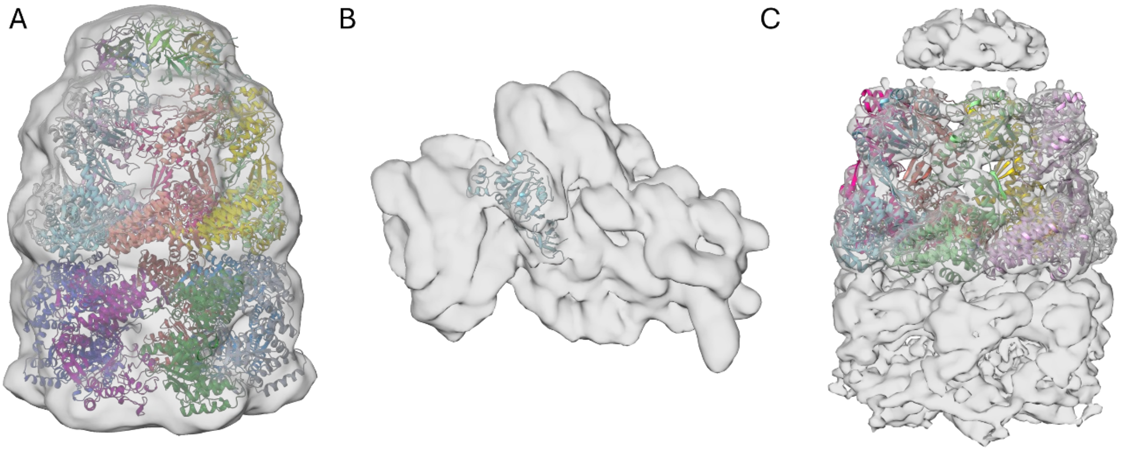
